# Extraordinary diversity of HLA class I gene expression in single cells contribute to the plasticity and adaptability of human immune system

**DOI:** 10.1101/725119

**Authors:** Rui Tian, Hao Zhu, Zhiying Pang, Yi Tian, Chao Liang

## Abstract

HLA, the coding genes of human major histocompatibility (MHC) proteins, play a crucial role in the human adaptive immune system by presenting antigenic peptides to T cell receptors on T cells. HLA-A, HLA-B and HLA-C, these 3 Class I HLA genes are one of the most polymorphic loci in the human genome. For decades, HLA typing has been performed prior to tissue and stem cell transplantation. However, beyond the role in tissue matching, HLA has also been implicated in a wide array of autoimmune diseases and HLA genotypes and expression levels are closely associated with cancer patients prognosis as recent studies have revealed. Recently methods have been developed to perform HLA typing and HLA expression quantification together by using RNA-seq techniques. However, these bulk RNA-seq experiments are measuring an averaged signal of cell populations. Single-cell RNA-seq (scRNA-seq) has regained its popularity due to its power to reliably resolve single RNA transcriptomes at large scales. In our present study, we did HLA typing using three independent scRNA-seq datasets. Interestingly, we found that single cells from the same donor could be classified into different groups where each group has a distinct expressed HLA genotype (e.g., HLA-A, heterozygous or homozygous); in other words, HLA class I genes show abundant allele specific expression in single cells. This phenomenon has been repeatedly observed in a total of 14 donors from 3 independent datasets (one is breast epithelium, another two are multiple myeloma). Our systematic analysis of HLA class I gene expression using multiple scRNA-seq datasets has uncovered a putative mechanism, where by fine tuning HLA class I expressions both at the quantity and allele levels, our immune system is able to handle various internal challenges through single cells equipped with extraordinary diverse HLA expression patterns.

## Introduction

HLA (human leukocyte antigen), the coding genes for MHC (human major histocompatibility), represent the core unit for the control of self versus non-self recognition by presenting antigenic peptides to the T cell receptor on T cells. HLA genes are mainly classified into two classes (although there is a 3^rd^ class which is not of primary research interests), HLA-A, HLA-B and HLA-C are the classic class I genes, while HLA-DR, HLA-DQ and HLA-DP are the most commonly mentioned class II genes. HLA is one of the most polymorphic loci in the human genome. The IMGT/HLA database released less than 1,000 HLA alleles (class I and II alleles combined) in 1998, this figure increased into 3,410 by the end of 2008 (the beginning of next generation sequencing age, NGS age); the latest figure is 23, 907 (up to 2019 July) and 17,191 are class I alleles accounting for 72% of all HLA alleles (https://www.ebi.ac.uk/ipd/imgt/hla/stats.html). NGS has brought almost a magnitude increase of newly registered HLA genotypes within ten years and also elevated the HLA typing to an unprecedented high resolution, shaking off those annoying mistyping and ambiguities[1–4].

Single-cell RNA-seq (scRNA-seq) represents a series of experimental techniques which allow us to quantify the transcriptomes of each individual cell at the scale of thousands of cells. With this robust technique, scientists could eventually investigate questions that have intrigued them for decades, such as the development of an early embryo, heterogeneity of stem cells and dynamics during stem cell differentiation, heterogeneity of cancer cells, tumor microenvironment, etc[5–13]. sc-RNA-seq could reveal cell to cell variability which could not be appreciated otherwise by traditional bulk RNA-seq, and cell population stratification or clustering is among the top-centered tasks.

In diploids, like human, two copies of genes from two parental sides do not necessarily express at the same level. This imbalance of allelic expressions could be as dramatic as mono-allelic expression (such as those imprinted genes or X chromosome deactivated genes), or more commonly seen as one parental allele expresses at a level that is significantly higher than another, referring to as allele specific expression (ASE)[14]. ASE, which is large attributable to cis-elements polymorphisms (i.e., sequence variations at promoter or enhancer regions), has been observed for more than half of SNP-distinguishable human genes and may contribute to human variability and heterosis in plants as well[14–17].

ASE is traditionally studied by sequencing tissues or cell populations (bulk RNA-seq), which measures the averaged signal of cell populations. scRNA-seq, however, allows us to get a picture of the expression of any given two alleles in each individual cell (provided the expression of this gene is detectable). HLA typing as mainly for the purpose of tissue or cell transplant matching are predominantly performed using DNA extracted from blood or buccal samples. In transplant practices, the importance of RNA or protein expression levels of HLA genes have been gradually realized. There are some reports on HLA typing using RNA-seq data and quantification of HLA genes as well[18–21]. However, to date, few reports have been available on the HLA typing at the single-cell level by using scRNA-seq data. In this study, we attempted to perform HLA typing using scRNA-seq data selected from public databases. We observed that single cells from the same donor could be classified into multiple groups where each group is characterized by a distinct HLA genotype (RNA level). Our results were repeatedly seen in multiple independent datasets, suggesting that single cell diversity and variability in HLA expression is far more enormous than we could learn from tissue or cell population bulk RNA-seq experiments. We conclude that extraordinary polymorphism at HLA class I alleles (DNA level) plus ultra-plasticity at RNA level per allele per single cell play an essential role in getting our immune system ready for dealing with unlimited internal and external challenges.

## Results and Discussion

Firstly we attempted to perform HLA genotyping using a scRNA-seq dataset derived from breast epithelium. As per cell sequencing depth varies and HLA genes expression may vary across cells, we only kept those cells where at least one HLA class I gene (HLA-A, HLA-B, HLA-C) was successfully genotyped. In the breast SRP140489 dataset[22], a total of 75 single cells were retained. Among these 75 cells, we could reliably report 13 cells with A*30:01/A*30:01, 12 cells with A*33:01/A*33:01 and 6 cells with A*30:01/A*33:01; 19 cells with B*14:02/B*14:02, 10 cells with B*13:02/B*13:02; 10 cells with C*08:02/C*08:02, 9 cells with C*06:02/C*06:02, 7 cells with C*06:02/C*08:02 (Table 1). To sum up those HLA-A, HLA-B and HLA-C genotyped cells separately, one comes up to 31, 29, 26, which is not equal to one another. The underlying reason is that, we seldom saw cells with all three HLA genes genotyped simultaneously, most often the case is that, you saw in a single cell one or two HLA genes genotyped, leaving two or one HLA genes undetectable due to insufficient reads coverage.

**Table 1.** HLA class I genes typing at RNA levels in single cells of multiple datasets.

| Datasets | Donors | HLA-A | Cell counts | HLA-B | Cell counts | HLA-C | Cell counts |
| --- | --- | --- | --- | --- | --- | --- | --- |
| SRP132719<br>Multiple myeloma (CD138+) | MM16 | A*24:02/A*24:02 | 12 | B*40:01/B*40:01 | 16 | C*03:03/C*03:03 | 11 |
|  |  |  |  |  |  | C*03:02/C*03:03 | 2 |
|  |  |  |  |  |  | C*03:02/C*03:02 | 2 |
|  | MM34 | A*24:02/A*24:02 | 70 | B*40:01/B*54:01 | 38 | C*01:02/C*03:04 | 36 |
|  |  |  |  | B*40:01/B*40:01 | 21 | C*03:04/C*03:04 | 28 |
|  |  |  |  | B*54:01/B*54:01 | 12 |  |  |
| SRP140489<br>breast epithilium | Donor1 | A*30:01/A*30:01 | 13 | B*14:02/B*14:02 | 19 | C*08:02/C*08:02 | 10 |
|  |  | A*33:01/A*33:01 | 12 | B*13:02/B*13:02 | 10 | C*06:02/C*06:02 | 9 |
|  |  | A*30:01/A*33:01 | 6 |  |  | C*06:02/C*08:02 | 7 |
| SRP158590<br>Multiple myeloma (CD138+) | IgM-MGUS1 | A*24:02/A*24:02 | 2 | B*18:01/B*56:01 | 4 | C*05:01/C*05:01 | 3 |
|  |  | A*24:02/A*32:01 | 1 | B*18:01/B*18:01 | 2 | C*01:02/C*05:01 | 2 |
|  |  | A*32:01/A*32:01 | 1 | B*56:01/B*56:01 | 2 | C*01:02/C*01:02 | 1 |
|  | NDMM3 | A*32:01/A*32:01 | 12 | B*18:01/B*51:01 | 17 | C*12:03/C*15:02 | 29 |
|  |  | A*25:01/A*25:01 | 10 | B*18:01/B*18:01 | 10 | C*15:02/C*15:02 | 8 |
|  |  | A*25:01/A*32:01 | 4 | B*51:01/B*51:01 | 9 | C*12:03/C*12:03 | 4 |
|  | NDMM6 | A*02:01/A*02:01 | 7 | B*44:03/B*44:03 | 4 | C*16:01/C*16:01 | 6 |
|  |  | A*02:01/A*29:02 | 2 | B*35:01/B*35:01 | 2 | C*04:01/C*04:01 | 5 |
|  |  | A*29:02/A*29:02 | 2 |  |  | C*04:01/C*16:01 | 4 |
|  | NDMM7 | A*25:01/A*25:01 | 6 | B*07:02/B*07:02 | 5 | C*06:02/C*06:02 | 10 |
|  |  |  |  | B*07:02/B*57:01 | 5 | C*06:02/C*07:02 | 1 |
|  |  |  |  | B*57:01/B*57:01 | 3 |  |  |
|  | NDMM8 | A*02:01/A*02:01 | 32 | B*41:02/B*41:02 | 17 | C*06:02/C*06:02 | 15 |
|  |  |  |  | B*13:02/B*41:02 | 8 | C*06:02/C*17:01 | 11 |
|  |  |  |  | B*13:02/B*13:02 | 6 | C*17:01/C*17:01 | 4 |
|  | RRMM1 | A*32:01/A*32:01 | 7 | B*44:02/B*44:02 | 8 | C*02:02/C*05:01 | 32 |
|  |  | A*24:02/A*24:02 | 5 | B*44:02/B*44:05 | 8 |  |  |
|  |  | A*24:02/A*32:01 | 2 | B*44:05/B*44:05 | 8 |  |  |
|  | RRMM2 | A*02:01/A*02:01 | 12 | B*44:02/B*44:02 | 8 | C*05:01/C*05:01 | 24 |
|  |  | A*02:01/A*03:01 | 2 | B*07:02/B*44:02 | 7 | C*05:01/C*07:02 | 6 |
|  |  | A*03:01/A*03:01 | 1 | B*07:02/B*07:02 | 6 |  |  |
|  | RRMM4 | A*02:01/A*02:01 | 6 | B*38:01/B*38:01 | 7 | C*12:03/C*12:03 | 6 |
|  |  | A*02:01/A*03:01 | 5 | B*35:01/B*38:01 | 3 | C*04:01/C*12:03 | 4 |
|  |  | A*03:01/A*03:01 | 4 |  |  | C*04:01/C*04:01 | 2 |
|  | SMM0 | A*03:01/A*03:01 | 14 | B*07:02/B*35:01 | 27 | C*04:01/C*04:01 | 35 |
|  |  | A*11:01/A*11:01 | 6 | B*07:02/B*07:02 | 14 | C*04:01/C*07:02 | 16 |
|  |  | A*03:01/A*11:01 | 3 | B*35:01/B*35:01 | 7 |  |  |
|  | SMM3 | A*01:01/A*01:01 | 10 | B*08:01/B*08:01 | 8 | C*07:01/C*07:01 | 9 |
|  |  |  |  | B*08:01/B*18:01 | 7 | C*03:14/C*07:01 | 1 |
|  |  |  |  | B*18:01/B*18:01 | 3 |  |  |
|  | SMM4 | A*02:01/A*02:01 | 7 | B*44:02/B*44:02 | 6 | C*05:01/C*06:02 | 16 |
|  |  | A*01:01/A*01:01 | 3 | B*57:01/B*57:01 | 6 | C*05:01/C*05:01 | 6 |
|  |  | A*01:01/A*02:01 | 2 | B*44:02/B*57:01 | 5 | C*06:02/C*06:02 | 2 |

So apparently we observed allele specific expressions (ASE) for all HLA class I genes (classic) in the breast scRNA-seq dataset. To figure out whether this observation is solely present in this single dataset, or rather represents a more general phenomenon, we set out to examine two more independent scRNA-seq datasets which were generated from multiple myeloma (MM) patients’ CD138 positive plasma cells separated from bone marrow and extramedullary sites. Two donors from the SRP132719 dataset [23] show ASEs of HLA class I genes, as shown in Table 1, for instance the donor MM34, for the HLA-B gene expression, there are 38 cells of heterozygous genotype (B*40:01/B*54:01), 21 cells of homozygous (B*40:01/B*40:01) and 12 cells of another homozygous genotype (B*54:01/B*54:01). For the HLA-C locus, 36 cells are heterozygous while 28 cells are homozygous. For the HLA-A locus, it seems that there is no polymorphism at all. In another multiple myeloma patients scRNA-seq dataset, a total 11 donors from dataset SRP158590 [24] also shows ASE of HLA class I genes (as shown in Table 1). For example, the donor NDMM3 shows ASE for all three HLA class I genes. The cells counted based on per gene genotypes are as the following, The HLA-A:12, 10, 4(A*32:01/A*32:01, A*25:01/A*25:01, A*25:01/A*32:01); The HLA-B: 17, 10, 9(B*18:01/B*51:01, B*18:01/B*18:01, B*51:01/B*51:01); The HLA-C: 29, 8, 4(C*12:03/C*15:02, C*15:02/C*15:02, C*12:03/C*12:03).

We have shown that ASE of HLA class I genes could be observed at the single-cell level in three independent datasets, pointing to a somewhat general phenomenon which has been under-appreciated at cell population or tissue levels, in which scenario, both genotyping or RNA level quantification would reflect the sum or average of multiple single cells. One might argue that, the HLA genotyping in the present study using these single-cell RNA-seq datasets, might be not precise analytically. Theoretically, if very single cell is indeed heterozygous at the RNA level, the reportedly homozygous genotypes might be the result of failing to report another allele which truly exists. In order to rule out this possibility which would make our discovery an artifact, we checked the sequencing reads coverage on HLA loci to make sure the genotyping in our report is reliable. Figure 1 shows an example of genotyping using scRNA-seq data from one single cell from dataset SRP132719 (MM). This cell was genotyped as HLA-A*24:02, HLA-B*54:01/40:01 and HLA-C*03:04/01:02, in total there are 1631 reads covering the HLA-A, HLA-B and HLA-C regions. As one could clearly see from the per position read coverage plots that, homozygous is indeed homozygous and heterzygosity has sufficient reads support for both alleles.

**Figure 1.**
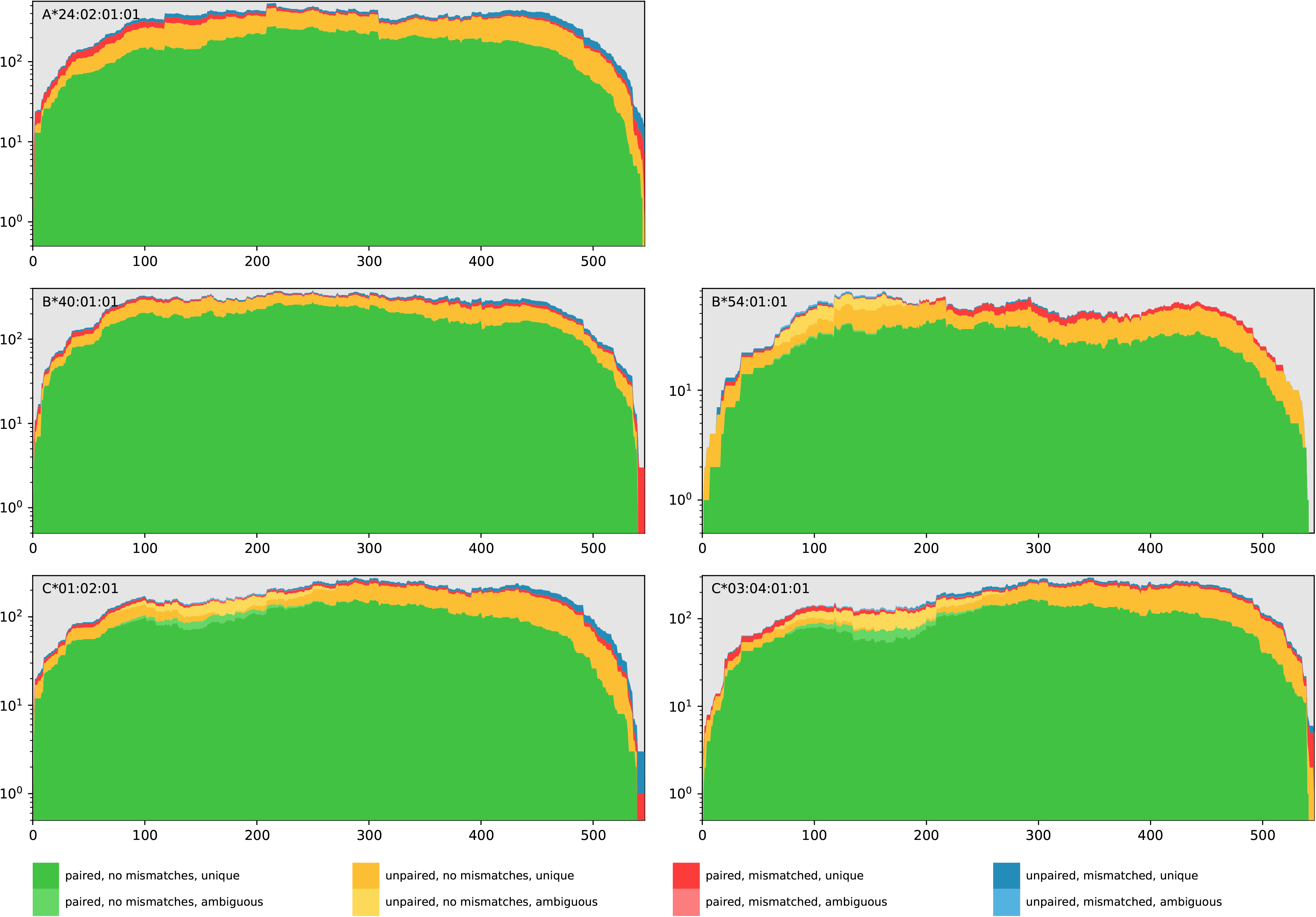
An example of a single cell HLA typing at the RNA level. Three rows represent the HLA-A, HLA-B and HLA-C locus respectively. This figure illustrates that at the RNA level, this single cell is of HLA-A homozygous (A*24:02), HLA-B heterozygous (B*40:01/B*54:01) and HLA-C heterozygous (C*01:01/C*03:04). For each panel, the x axis represents the concatenated exon 2 and exon 3 of HLA alleles, while the y axis denotes the reads counts fall onto each genomic position. This figure is intended to show as a proof that the HLA typing in the present study is reliable and accurate as it is supported with sufficient raw reads.

During the process of HLA genotyping of those single cells from three datasets (a total of 14 donors), we noticed that the HLA class I genes expression is highly imbalanced in a single cell, as such we were hardly able to genotype all three HLA class I genes simultaneously. Therefore, we visualized the expression levels of HLA-A, HLA-B and HLA-C genes in all the single cells per donor. As shown in Figure 2, the heatmap of HLA Class I genes explicitly exhibit the trend that in almost very single cell, there is at least one HLA class I gene is lowly expressed (or expression level undetectable). This observation also holds for multiple donors from multiple myeloma datasets. It is noteworthy that, almost all single cells from one of MM patient (6) are characterized by very low level of HLA-A expression (Figure 2). This might suggest an immune escape strategy by MM via repressing the expression of HLA-A antigens, although more efforts are needed before a definitive conclusion could be reached.

**Figure 2.**
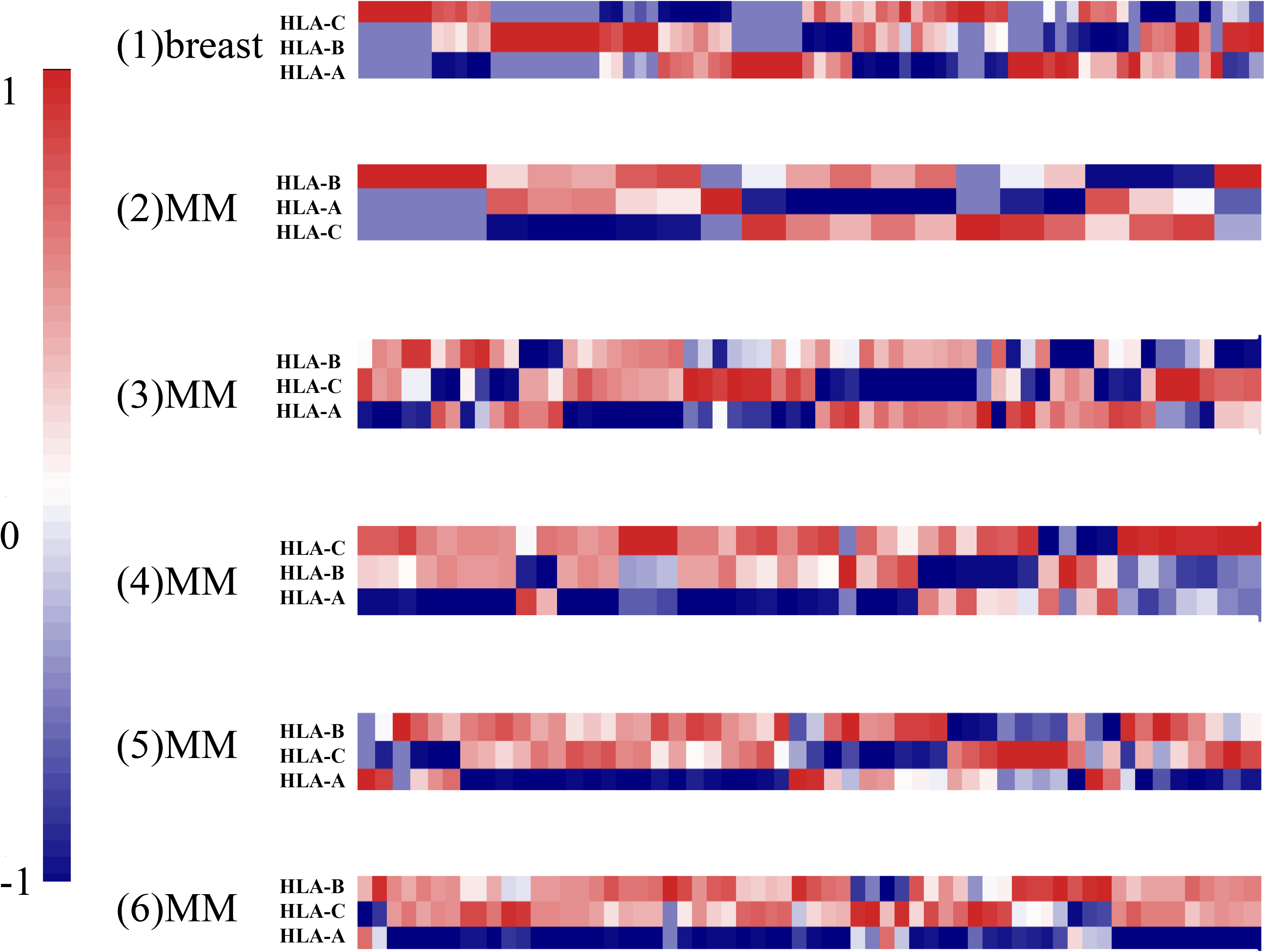
The heatmap of HLA-A, HLA-B, HLA-C expression in single cells across multiple donors (in total 6 donors are shown here). The first one is from breast epithelium and the other five are from multiple myeloma (MM) patients. The HLA class I genes expression levels were normalized and scaled by column before they were extracted separately for visualization purpose from the whole expression sheet encompassing all expressed genes per single cell. Heterogeneity of HLA class I genes expression could easily classify single cells into multiple different groups. It seems that in each single group, there is at least one HLA gene lowly expressed. For donor (6), a MM patient, it seems that the HLA-A is hardly detectable in vast majority of single cells.

In summary, we report ASE and imbalanced expression of HLA class I genes the single-cell level, we have provided compelling evidence from multiple independent datasets to support our discovery. Our findings in this study unravels the abundant variability and diversity of HLA class I genes expression at the single-cell level which is under-appreciated before. Combined with other mechanisms, we believe that this diversity and variability ultimately could enhance the immune adaptation for the survival of human beings under harsh and challenging environments.

## Material and Methods

Three independent scRNA-seq datasets were downloaded from NCBI GEO databases, namely two multiple myeloma (MM) scRNA-seq datasets (SRA study numbers: SRP132719, SRP158590) and one breast epithelium single cell dataset (SRA study number: SRP140489). These three datasets were chosen due to availability of raw data with sufficient sequencing depth per single cell. One of our general observations is that, most 10Xgenomics single cell RNA-seq data, as they tend to sort several thousand, even more than 10,000 cells per experiment, given ordinary around 100 G bases sequencing amount, per cell sequencing depth is not adequate for reliable HLA genotyping.

Raw data was downloaded using NCBI sra-toolkit “prefetch” command, and transformed into fastq files using fastq-dump. Two software were used for HLA class I genotyping, namely seq2HLA[18] and OptiType[25]. In most cases the typing results from the two tools would be identical, in cases where two results did not agree with each other, manual examination was needed to rule out erroneous typing based on the principle that, regardless the number of single cells, there should not be more than 3 genotypes per HLA locus identified in one single donor (we did observe some erroneously typed single cells most likely due to insufficient reads coverage over the HLA region).

The HLA class I genes quantification was done using seq2HLA. For scRNA-seq transcriptome quantification, reads were aligned to reference genome (hg19) using STAR[26], per gene (transcript) reads were counted with HTSeq-count[27] and finally the expression level of each gene as measured by RPKM was calculated with in house scripts. Normalization was done by each donor where HLA genes RPKM together with other genes RPKM in each single cell from the same donor were normalized using quantile normalization method[28–30]. Then the normalized HLA genes RPKMs were eventually used for visualization of expression levels (via heatmap plotting function of R language) across different single cells from each single donor.

## Abbreviations

HLA: human leukocyte antigen
scRNA-seq: single-cell RNA-seq
ASE: allele specific expression

